# Dynamic changes in cerebral and peripheral markers of glutamatergic signaling across the human sleep-wake cycle

**DOI:** 10.1101/458885

**Authors:** Susanne Weigend, Sebastian C. Holst, Valérie Treyer, Ruth L. O’Gorman Tuura, Josefine Meier, Simon M. Ametamey, Alfred Buck, Hans-Peter Landolt

**Affiliations:** Psychopharmacology and Sleep Research, Institute of Pharmacology and Toxicology, University of Zürich, Zürich, Switzerland; Sleep & Health Zürich, University Center of Competence, University of Zürich, Zürich Switzerland; Department of Nuclear Medicine, University Hospital Zurich, Zürich, Switzerland; Institute for Regenerative Medicine (IREM), University of Zürich, Switzerland; Center of MR Research, Children’s University Hospital, Zürich, Switzerland; Radiopharmaceutical Science, Department of Chemistry and Applied Biosciences, Institute of Pharmaceutical Sciences, ETH Zurich, Zürich, Switzerland

**Keywords:** PET-MRS imaging, sleep homeostasis, FMRP, BDNF, plasticity

## Abstract

Both sleep and glutamatergic signaling in the brain are tightly controlled and homeostatically regulated. Sleep homeostasis is reliably reflected by predictable changes in brain electrical activity in waking and sleep, yet the underlying molecular mechanisms remain elusive. Current hypotheses posit that recovery sleep following prolonged waking restores efficient functioning of the brain, for example by keeping glutamatergic signaling in a homeostatic range. We recently provided evidence in humans and mice that metabotropic glutamate receptors of subtype-5 (mGluR5) contribute to the brain’s coping mechanisms with sleep deprivation. Here we combined in 31 healthy men, proton magnetic resonance spectroscopy to measure the levels of glutamate (Glu), GLX (glutamate-to-glutamine ratio) and GABA (γ-amino-butyric-acid) in basal ganglia (BG) and dorsolateral prefrontal cortex, simultaneous positron emission tomography to quantify mGluR5 availability with the novel radioligand, [^18^F]PSS232, and quantification in blood plasma of the mGluR5-regulated proteins, fragile-X mental retardation protein (FMRP) and brain-derived neurotrophic factor (BDNF). All measurements were conducted at the same circadian time in baseline, following sleep deprivation and after recovery sleep. We found that Glu and GLX in BG (p_all_ < 0.01), but not in prefrontal cortex, and the plasma concentration of FMRP (p < 0.02), were increased after sleep loss and tended to normalize following recovery sleep (p_all_ < 0.1). Furthermore, a night without sleep enhanced whole-brain and striatal mGluR5 availability and was normalized by recovery sleep (p_all_ < 0.05). By contrast, other brain metabolites and plasma BDNF levels were not altered. The findings demonstrate convergent changes in distinct markers of glutamatergic signaling across prolonged wakefulness and recovery sleep in humans. They warrant further studies to elucidate the underlying mechanisms that link the homeostatic regulation of sleep and glutamatergic system activity in health and disease.

**One-sentence summary:** Sleep-dependent recovery of wakefulness-induced changes in, cerebral glutamatergic signaling

**Major subject area:** Neuroscience; Human Biology & Medicine

## Introduction

Sleep has been conserved throughout evolution and is generally assumed to fulfill vital biological functions. This notion is corroborated by the general principle referred to as sleep homeostasis, which assumes that the lack of sleep is predictably compensated by increased sleep need and intensity as reflected by electroencephalographic (EEG) slow-wave activity (SWA; activity in the ~ 0.75-4.5 Hz range) in non-rapid-eye-movement NREM sleep (Achermann and Borbély, 2017). Prevailing current hypotheses posit that sleep homeostasis serves the normalization of synaptic long-term potentiation (LTP) occurring during wakefulness, by synaptic long-term depression (LTD) occurring during NREM sleep (Tadavarty et al., 2009; Pigeat et al., 2015; Tononi & Cirelli, 2014).

Glutamate (Glu) plays an essential role in the fine-tuned molecular processes underpinning LTP and LTD (Huber et al., 2002; Marshall, 2006, Kauer and Malenka, 2007). Overstimulation of metabotropic and ionotropic Glu receptors by excess extracellular Glu is a major culprit of neuronal excitotoxicity and contributes to neuropsychiatric and neurodevelopmental disorders that can be exacerbated by inadequate sleep (Sanacora at al., 2008; Ahmed et al., 2011; Averill et al., 2017). Suggesting an important contribution of glutamatergic signaling to sleep homeostasis and a role for sleep in keeping extracellular Glu in a homeostatic range, Glu levels in the frontal cortex of freely moving rats rose during prolonged wakefulness and REM sleep and decreased during NREM sleep (Dash et al., 2009). No comparable data are currently available in humans.

Nevertheless, two key players were recently identified that may orchestrate synaptic plasticity and glutamatergic signaling across the sleep-wake cycle: Homer1a and metabotropic Glu receptors of subtype-5 (mGluR5). Homer1a uncouples mGluR5 from their downstream signaling partners, which leads to synaptic LTD (Kammermeier and Worley, 2007; Berridge, 2016; Ronesi and Huber, 2008). Biochemical, proteomic and imaging studies in mice demonstrated that Homer1a and signaling from group-I mGluRs (mGluR1/5) drive the homeostatic downscaling of excitatory synapses during sleep (Diering et al., 2017). In humans, mGluR5 show high expression in brain regions regulating sleep (Hefti et al., 2013) and their functional availability was increased after prolonged wakefulness (Hefti et al., 2013). Furthermore, increased mGluR5 availability correlated with behavioral and neurophysiological markers of elevated sleep need, including self-rated sleepiness, unintended sleep during prolonged wakefulness, as well as SWA and slow (< 1 Hz) oscillatory activity in the NREM sleep EEG (Hefti et al., 2013; Holst et al., 2017).

Apart from interacting with Homer1a, activation of mGluR5 regulates the expression of fragile X mental retardation protein (FMRP) and brain-derived neurotrophic factor (BDNF), which both play important roles in neuronal plasticity (Comery et al., 1997; Huber et al., 2002; Li et al., 2002; Restivo et al., 2005; Bramham and Messaoudi, 2005; Desai et al., 2006; Lu et al., 2014). Work in Drosophila suggested that the *dFmr1* gene is a molecular regulator of sleep need (Bushey et al., 2009) and that the expression of FMRP controls sleep time and the sleep loss-induced sleep rebound (Bushey et al., 2011). Similarly, the expression of BDNF protein in mice has been associated with the rebound in SWA following sleep deprivation (Huber et al., 2007). Whereas the effects of prolonged waking on the concentration of FMRP in humans are unknown, for BDNF either an increase (Schmitt et al., 2016) or a decrease (Kuhn et al., 2016) have been reported.

Based upon the evidence outlined above, we simultaneously quantified in healthy human volunteers dynamic changes in brain metabolites, including GLX, Glu and GABA (γ-amino-butyric-acid) in dorso-lateral prefrontal cortex (dlPFC) and basal ganglia (BG), cerebral mGluR5 availability, as well as FMRP and BDNF levels in blood serum after prolonged wakefulness and following recovery sleep. We hypothesized that sleep loss increases these potential markers of elevated sleep need and expected that recovery sleep normalizes the waking induced changes. With the exception of BDNF, all markers quantified revealed the expected changes, suggesting that glutamatergic signaling involving mGluR5 importantly contributes to the regulation of sleep-wake dependent synaptic plasticity in humans.

## Results

Thirty-one healthy men completed this strictly controlled study (Table 1 for demographics; the numbers of study participants contributing to each analysis are specified below). Following 8-hour adaptation and baseline sleep opportunities in the sleep laboratory, all volunteers stayed awake under constant supervision for 40 hours, followed by a 10-hour recovery sleep opportunity. All measurements in baseline (BL), after sleep deprivation (SD) and after recovery (RE) sleep were conducted at the same circadian time in all three conditions, starting at 4:23 pm ± 23 min (Fig. 1).

**Table 1.**
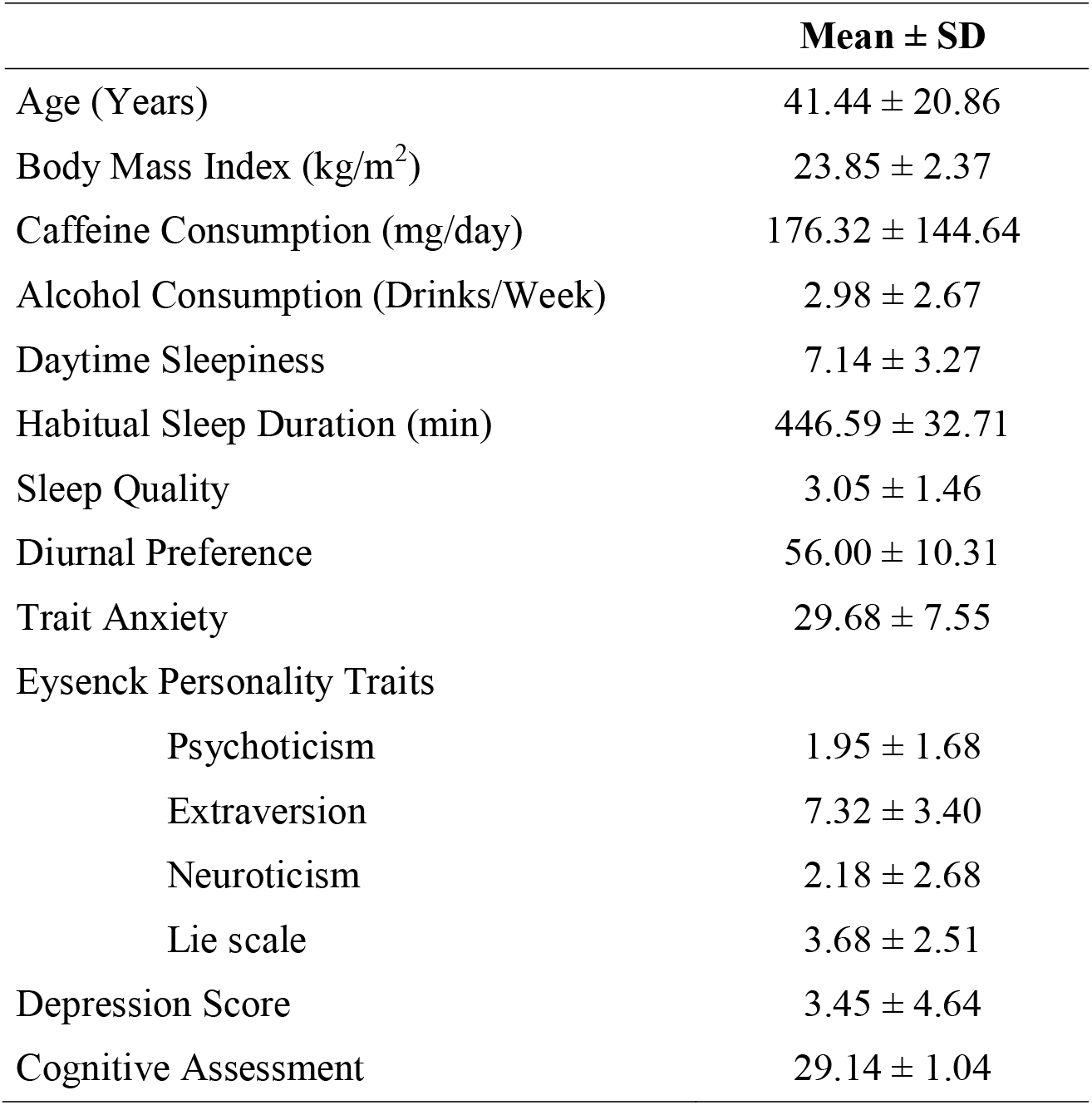
Demographic Data of Study Participants. Values represent means ± SEM (n = 31). Caffeine consumption was calculated based on the following amounts per serving: coffee: 100mg; ceylon or green tea: 30 mg; cola drink: 40 mg (2 dL); energy drink: 80 mg (2 dL); chocolate: 50 mg (100 g). Diurnal preference: Horne-Östberg Morningsness-Eveningness Questionnaire (Horne et al., 1976); daytime sleepiness: Epworth Sleepiness Scale (Bloch et al., 1999); depression score: Beck Depression Inventory (Beck et al., 1961); personality traits: Eysenck Personality Questionnaire (Francis et al., 2006); cognitive assessment: Montreal Cognitive Assessment (Nasreddine, 2005); trait anxiety: State-Trait Anxiety Inventory (Spielberger et al., 1970); sleep quality: Pittsburgh Sleep Quality Index (Buysse et al., 1989).

|  | **Mean ± SD** |
| --- | --- |
| Age (Years) | 41.44 ± 20.86 |
| Body Mass Index (kg/m<sup>2</sup>) | 23.85 ± 2.37 |
| Caffeine Consumption (mg/day) | 176.32 ± 144.64 |
| Alcohol Consumption (Drinks/Week) | 2.98 ± 2.67 |
| Daytime Sleepiness | 7.14 ± 3.27 |
| Habitual Sleep Duration (min) | 446.59 ± 32.71 |
| Sleep Quality | 3.05 ± 1.46 |
| Diurnal Preference | 56.00 ± 10.31 |
| Trait Anxiety | 29.68 ± 7.55 |
| Eysenck Personality Traits |  |
| Psychoticism | 1.95 ± 1.68 |
| Extraversion | 7.32 ± 3.40 |
| Neuroticism | 2.18 ± 2.68 |
| Lie scale | 3.68 ± 2.51 |
| Depression Score | 3.45 ± 4.64 |
| Cognitive Assessment | 29.14 ± 1.04 |

**Figure 1.**
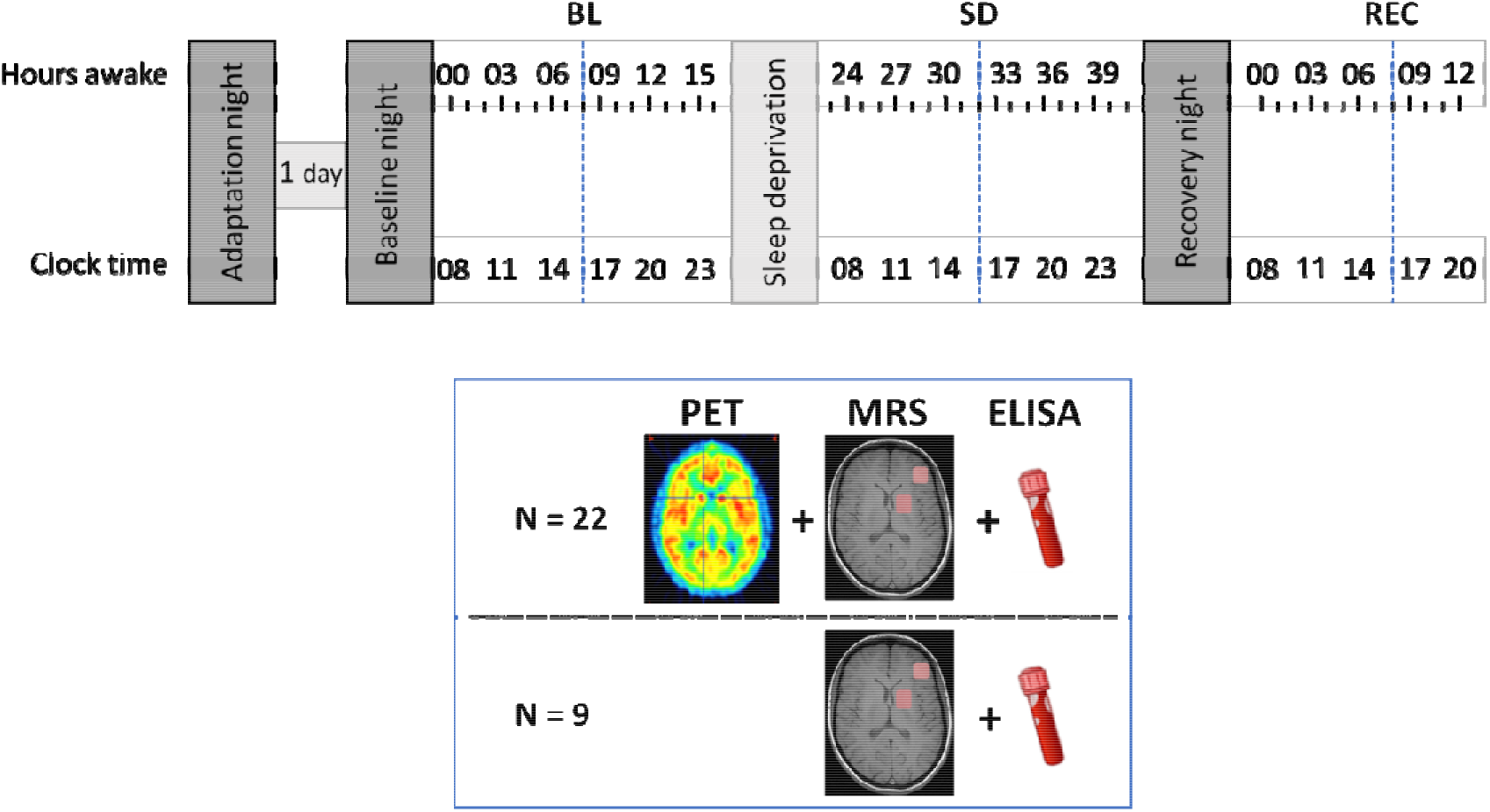
Experimental protocol. After an adaptation and baseline night, subjects underwent 40hrs of prolonged wakefulness followed by a recovery night. At baseline (BL), after sleep deprivation (SD), and again after recovery sleep (REC), levels of mGluR5 were measured using positron emission tomography with [^18^F]PSS232 at the same circadian timepoint (blue dotted lines). Furthermore, distinct brain metabolites were measured with magnetic resonance spectroscopy and blood samples for the quantification of blood BDNF and FMRP levels were drawn at these timepoints. Blue box summarizes type of data collection and number of subjects at the imaging sessions in BL, SD and REC conditions (blue dotted lines). A cognitive test session was performed every three hours of wakefulness consisting of vigilance (Psychomotor Vigilance Task (PVT); Dinges et al., 1985), sleepiness (Karolinska Sleepiness Scale (KSS); Akerstedt et al., 1990), tiredness symptoms (Tiredness Symptoms Scale (TSS); Schulz et al., 1991) and affective state (Visual Analogue Scales (VAS); Hoddes et al., 1973) testing.

### Sleep deprivation increases Glu and GLX levels in the basal ganglia

Methodological advances in proton magnetic resonance spectroscopy (H^1^-MRS) have recently permitted the non-invasive detection of naturally occurring changes in tightly regulated metabolite concentrations in circumscribed areas of the human brain. Whereas one recent study suggested that GLX levels in the left parietal lobe decrease over night (Volk et al., 2018), our own research revealed no significant changes after sleep deprivation in GLX/Glu and GABA in the medial prefrontal cortex (Holst et al., 2017). Thus, the exact roles in humans of the main excitatory and inhibitory neurotransmitters in circadian and homeostatic sleep-wake regulation remain unclear. Here, we quantified at the same circadian time in 20 study participants the effects of prolonged wakefulness and recovery sleep on the extracellular concentrations of Glu, GLX and GABA in two predefined voxels located in the cortex (dlPFC) and the BG. Both these regions show pronounced waking-induced changes in mGluR5 availability (Holst et al., 2017) and are thought to contribute importantly to sleep homeostasis (Dahan et al., 2006; Léna et al., 2005; Guillaumin et al., 2018). Consistent with our previous study (Holst et al., 2017), sleep deprivation caused no reliable changes in these metabolites in the cortex (Fig. 2, left-hand panel). By contrast, Glu and GLX levels in the BG were increased after prolonged waking in 17 of 20 study participants when compared to baseline (Fig. 2, right-hand panel). The mean increase in Glu equaled 6.3 ± 2.06 % (BL: 1.41 ± 0.02 [arbitrary units]; SD: 1.50 ± 0.03; SD *vs.* BL: p < 0.02, Tukey’s test, n = 20). Similarly, sleep loss increased the GLX concentration in the BG in 16 of 20 subjects, and the mean increase equaled 9.0 ± 2.53 % (BL: 1.66 ± 0.04; SD: 1.81 ± 0.05; SD *vs.* BL: p < 0.004). Although both, Glu (SD: 1.50 ± 0.03; RE: 1.45 ± 0.03; RE vs. SD: 2.8 % reduction) and GLX (SD: 1.81 ± 0.05; RE: 1.73 ± 0.04; RE *vs.* SD: 4.2 % reduction) were slightly reduced after recovery sleep when compared to sleep deprivation, these changes did not reach statistical significance.

**Figure 2:**
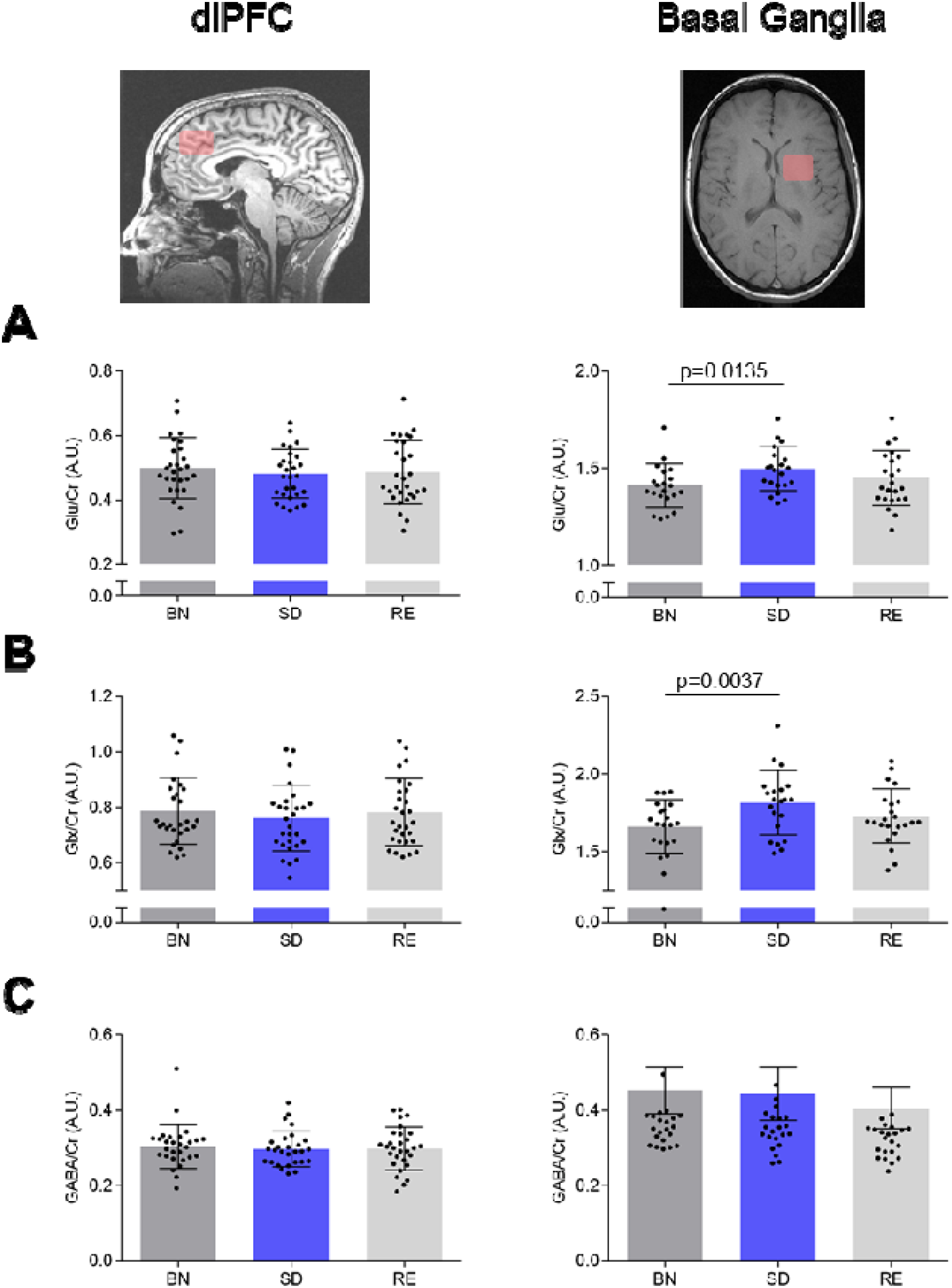
Effects of sleep deprivation and recovery sleep on endogenous brain metabolites in dorsolateral prefrontal cortex (left) and basal ganglia (right). Magnetic resonance spectroscopy yielded levels of glutamate (Glu; A), glutamate/glutamine ratio (Glx; B) and γ-aminobutyric acid (GABA; C) relative to creatine in baseline (BL, dark grey), sleep deprivation (SD, blue) and recovery (RE, light grey) conditions. Data represent means of arbitrary units (A.U.) ± standard error of the mean (SEM) in N=31. Black dots represent individual subjects. Data for Glu and Glx were acquired with PRESS and data for GABA with MEGAPRESS sequences. BG: Mixed-model ANOVA with factor ‘condition’: Glu - F_2,36_ = 4.83, p < 0.05; GLX - F_2,36_= 6.32, p < 0.01; GABA - F_2,36_ = 4.71, p < 0.02.

The levels of GABA remained stable in the BG following sleep deprivation and recovery sleep (BL: 0.45 ± 0.01; SD: 0.45 ± 0.007; SD vs. BL: p >0.8; SD: 0.45 ± 0.007; RE: 0.41 ± 0.01; p > 0.05, Tukey’s test, n = 20) (Fig. 2C). Similarly, no significant changes in other metabolites (N-acetylaspartate, glutathione, choline) were detected.

### Whole-brain mGluR5 availability is elevated after sleep deprivation and normalized after recovery sleep

To quantify sleep-wake associated changes in the availability of mGluR5 that may occur simultaneously with the above described local changes in Glu, GLX and GABA, the newly developed, highly selective, non-competitive mGluR5 antagonist for PET brain imaging, [^18^F]PSS232, was employed (Sephton et al., 2014; Warnock et al., 2018).

When compared to baseline, sleep deprivation induced a consistent increase in whole-brain [^18^F]PSS232 binding potential reflecting elevated cerebral mGluR5 availability (BL: 1.16 ± 0.04; SD:1.20 ± 0.04; SD *vs.* BL: p < 0.05, Tukey’s test) (Fig. 3). The [^18^F]PSS232 binding increased from BL to SD in 15 of 20 subjects in whom PET scans in both conditions were available. On average, the sleep deprivation-induced increase in whole-brain mGluR5 availability equaled 5.53± 2.22 %.

**Figure 3:**
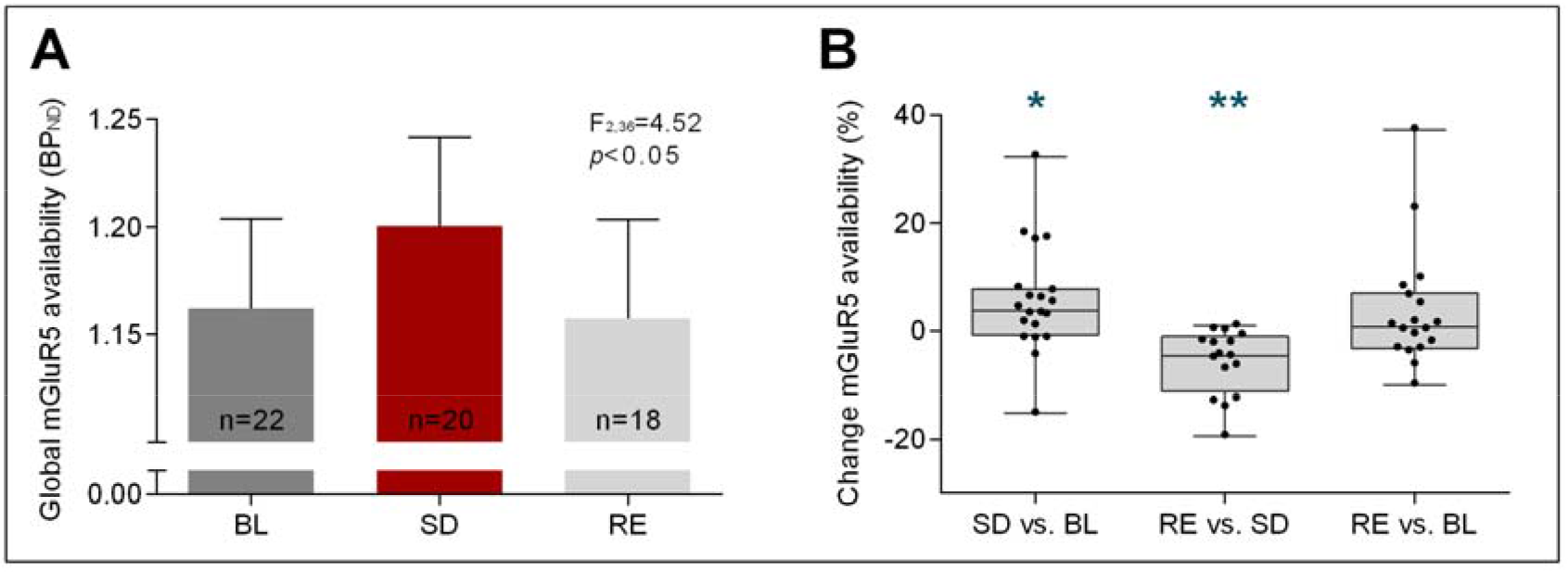
Effects of sleep deprivation and recovery sleep on metabotropic glutamate receptor subtype 5 availability. (A) Global NonDisplaceable binding potential (BPND) after [^18^F]PSS232 uptake in human brain. Columns display global mGluR5 availability in the human brain in baseline (BL, dark grey), sleep deprivation (SD, red) and recovery (RE, light grey) conditions. Data represent means + standard error of the mean (SEM) in N=22 in baseline, N=20 in sleep deprivation and N=18 in recovery condition. (mixed-model ANOVA, factor ‘condition’: F_2,36_=4.52, p<0.05) (B) Box plots illustrate the calculated increase of global mGluR5 availability in percent after sleep deprivation (SD vs. BL), the calculated decrease after recovery sleep (RE vs. SD) and in addition the difference between baseline and recovery conditions (RE vs. BL). Black dots represent individual subjects. Asteriks indicate significant increase or decrease in change scores (Mann-Whitney *U* tests: * = p < 0.05; ** = p < 0.01).

To examine whether recovery sleep reverses the wakefulness-induced changes, PET scans were also performed after the recovery night. In 13 of 16 study participants in whom SD and RE data were available, whole-brain [^18^F]PSS232 binding was reduced in RE when compared to SD (SD: 1.21 ± 0.05; RE: 1.14 ± 0.04; RE *vs.* SD: p < 0.01, Tukey’s test). The reduction in mGluR5 availability from SD to RE equaled 5.77± 1.50 %. No difference in [^18^F]PSS232 binding potential between BL and RE was detected, suggesting that recovery sleep normalized the waking-induced enhancement in mGluR5 availability.

### Wake-sleep dependent changes in mGluR5 availability in the basal ganglia

Given the waking-induced increase in Glu and GLX in the BG and the fact that the striatum and the amygdala show high mGluR5 expression (Gasparini et al., 2008; Hefti et al., 2013), the wake-sleep associated changes in [^18^F]PSS232 binding were quantified specifically in caudate nucleus, putamen and amygdala. A pronounced increase in mGluR5 availability after prolonged waking was confirmed in all three regions (caudate nucleus: BL: 1.15 ± 0.06; SD: 1.25 ± 0.06; increase: 8.7 ± 4.8 %; SD *vs.* BL: p < 0.03; putamen: BL: 1.18 ± 0.05; SD: 1.20 ± 0.05; increase: 5.41 ± 2.42 %; SD *vs.* BL: p < 0.2; amygdala: BL: 1.27 ± 0.07; SD: 1.38 ± 0.07; increase: 8.6 ± 4.72 %; SD *vs.* BL: p < 0.03, Tukey’s tests, n = 20) (Fig. 4). Similar to the whole-brain data, recovery sleep normalized mGluR5 availability in caudate nucleus (SD: 1.25 ± 0.06; RE: 1.14 ± 0.06; reduction: 8.59 ± 3.46 %; RE *vs.* SD: p < 0.02), putamen (SD: 1.23 ± 0.05; RE: 1.16 ± 0.05; reduction: 6.09 ± 2.65 %; RE *vs.* SD: p < 0.02) and amygdala (SD: 1.38 ± 0.07; RE: 1.23 ± 0.07; reduction: 11.31 ± 4.71 %; RE vs. SD: p < 0.01, Tukey’s tests, n = 16) to the level of baseline (RE *vs.* BL: p_all_ > 0.5, Tukey’s test, n = 16) (Fig. 4).

**Figure 4:**
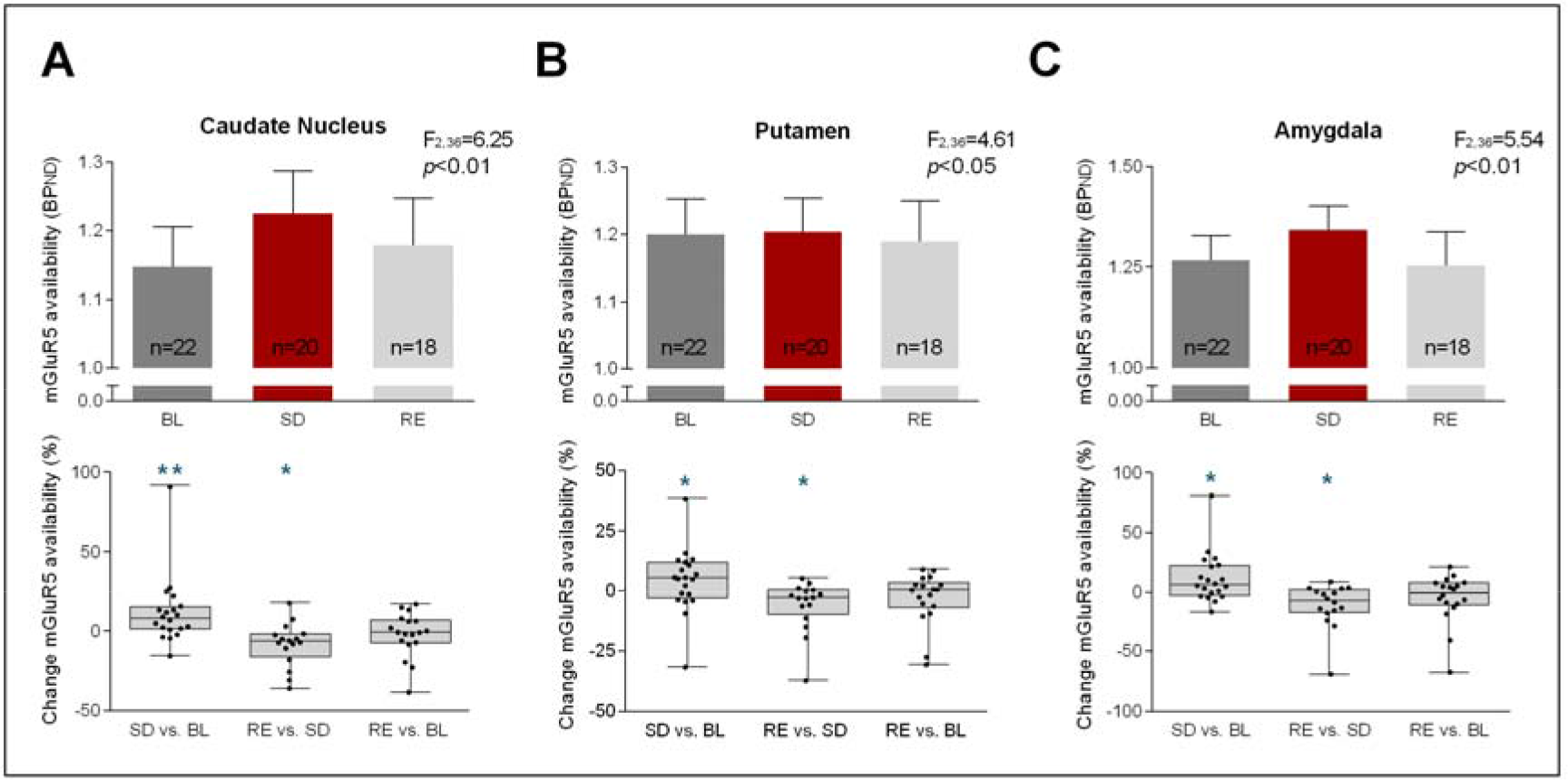
Regional differences in the effect of sleep deprivation and recovery sleep on metabotropic glutamate receptor subtype 5. Top: NonDisplaceable binding potential (BP_ND_) after [^18^F]PSS232 uptake in Caudate Nucleus **(A)**, Putamen **(B)** and Amygdala **(C)**. Columns display global mGluR5 availability in the human brain in baseline (BL, dark grey), sleep deprivation (SD, red) and recovery (RE, light grey) conditions. Data represent means + standard error of the mean (SEM) in N=22 in baseline, N=20 in sleep deprivation and N=18 in recovery condition. (mixed-model ANOVA, factor ‘condition’ Caudate Nucleus: F_2,36_= 6.25, p<0.01; Putamen: F_2,36_=4.61, p<0.05; Amygdala: F_2,36_=5.54, p<0.01) Bottom: Box plots illustrate the calculated increase of mGluR5 availability in percent in Caudate Nucleus **(A)**, Putamen **(B)** and Amygdala **(C)** after sleep deprivation (SD vs. BL), the calculated decrease after recovery sleep (RE vs. SD) and in addition the difference between baseline and recovery conditions (RE vs. BL). Black dots represent individual subjects. Asteriks indicate significant increase and decrease in change scores (Mann-Whitney *U* tests: * = p < 0.05; ** = p < 0.01).

### Sleep deprivation increases FMRP concentration in blood serum

To tackle the question whether the wake-sleep-related changes in Glu/GLX concentrations and mGluR5 availability in the brain are mimicked by changes in mGluR5-regulated proteins in peripheral blood, circulating FMRP and BDNF in serum were quantified with enzyme-linked immunosorbent assays (ELISA) in BL, SD and RE conditions. Intriguingly, prolonged waking increased blood FMRP concentration by 25.86 ± 16.39 % (BL: 268.52 ± 33.76 pg/ml; SD: 370.86 ± 31.93 pg/ml; SD *vs.* BL: p < 0.02, n = 23) (Fig. 5). Although the FMRP concentration tended to revert to baseline, the values in SD and RE conditions were not significantly different (SD: 370.86 ± 31.93 pg/ml; SD: 333.89 ± 33.51 pg/ml; RE *vs.* SD, p > 0.6). In contrast to FMRP, the levels of BDNF were not affected by prolonged waking nor recovery sleep (supplementary Figure S2).

**Figure 5:**
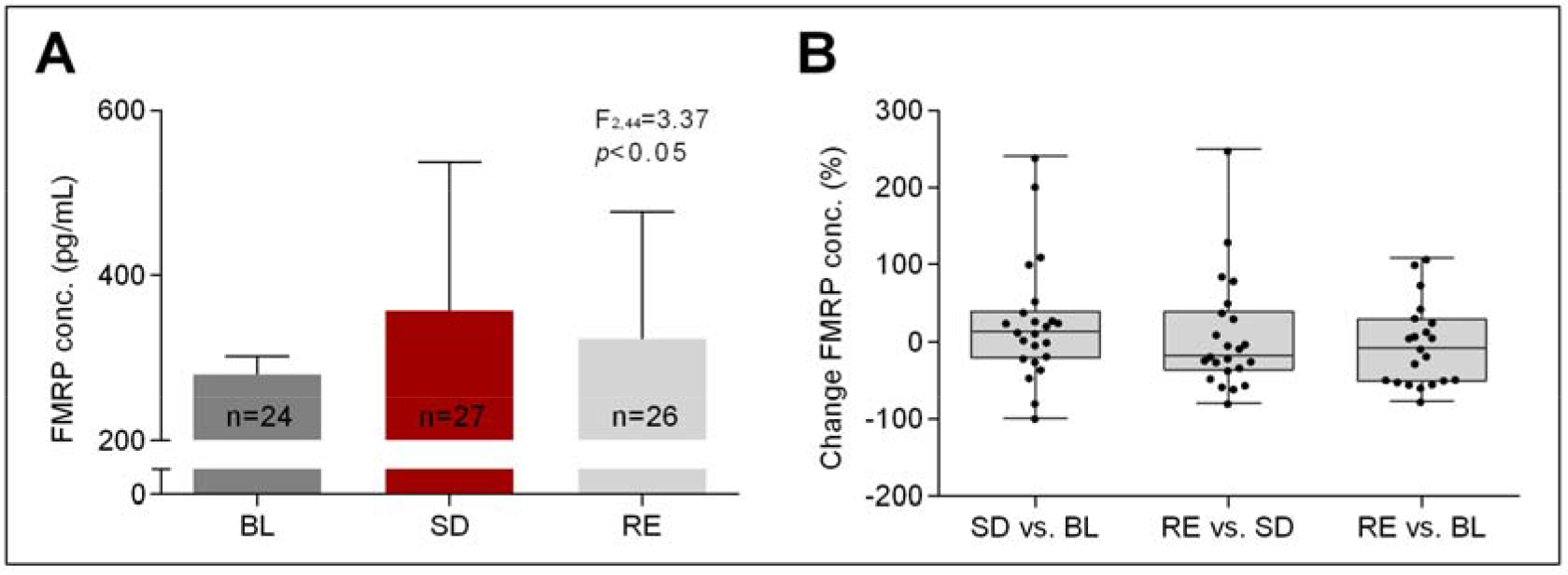
Effects of sleep deprivation and recovery sleep on blood fragile X mental retardation protein levels. **(A)** Circulating human blood levels of FMRP in pg/mL. Columns display amounts of FMRP in baseline (BL, dark grey), sleep deprivation (SD, red) and recovery (RE, light grey) conditions. Data represent means + standard error of the mean (SEM) in N=24 in baseline, N=27 in sleep deprivation and N=26 in recovery condition. Mixed model ANOVA main effect of ‘condition’: F_2, 44_ = 3.37, p < 0.05. (B) Box plots illustrate the calculated increase of blood FMRP levels in percent after sleep deprivation (SD vs. BL), the calculated decrease after recovery sleep (RE vs. SD) and in addition the difference between baseline and recovery conditions (RE vs. BL). Black dots represent individual subjects.

## Discussion

Glutamate is the primary excitatory neurotransmitter of the human brain. Although basic research *in vitro* and in animal models highlights a prominent role for glutamatergic mechanisms in regulating sleep-wake homeostasis (Maret et al., 2007; Dash et al., 2009; Ahnaou et al., 2015; Diering et al., 2017; Holst et al, 2017; for review, see Halassa & Haydon, 2010), knowledge about glutamatergic signaling as a function of waking and sleep in humans is scarce. Here we investigated the effects of prolonged wakefulness and recovery sleep on simultaneous changes in upstream and downstream molecular markers of metabotropic glutamatergic neurotransmission in healthy adults. We found that one night without sleep elicited reliable increases in cerebral Glu/GLX levels and mGluR5 availability, particularly in the basal ganglia, as well as in the concentration of the mGluR5-regulated protein, FMRP, in the blood stream. Given that these wakefulness-induced molecular changes tended to normalize after recovery sleep, the findings suggest that sleep is essential to keep glutamatergic signaling in a homeostatic range. Furthermore, the study indicates that human sleep may counteract neuronal dysfunction and degeneration, which can be caused by excessive glutamate (Sanacora at al., 2008; Ahmed et al., 2011; Averill et al., 2017), on multiple levels of the metabotropic glutamatergic signaling cascade.

### Sleep deprivation and recovery sleep induce dynamic changes in basal ganglia glutamate levels

The levels of glutamate in the rat cortical extra-synaptic space rise during waking and decrease during NREM sleep (Dash et al., 2009), yet it is currently unknown whether similar changes also occur in the human brain. To examine a glutamatergic contribution to the relief of depressive symptoms after wake therapy, brain levels of Glu, GLX and GABA were previously measured with ^1^H-MRS in depressed patients undergoing acute and repeated therapeutic sleep deprivation (Murck et al., 2002; Murck et al., 2009; Benedetti et al., 2009). No significant alterations in GLX or its elements were found in different cortical regions (dlPFC, anterior cingulate cortex and parieto-occipital cortex), yet preliminary data indicated that sleep loss increased GLX in subcortical brain regions (Murck et al., 2002). Because the baseline levels of GLX and Glu in cerebral cortex differ between depressed patients and healthy controls (Järnum et al., 2011; Njau et al., 2017), it is unclear whether these older studies are directly comparable with the present investigation. Nevertheless, our previous (Holst et al., 2017) and current work in healthy controls is consistent with the data in depressed patients (Benedetti et al., 2009; Murck et al., 2009). It indicates that prolonged wakefulness does not reliably alter the MRS signal compatible with GLX and its constituents in anterior cingulate cortex and dlPFC. It cannot be excluded, however, the lack of a significant change in GLX in the dlPFC voxel could be related to the voxel composition, which, compared to the basal ganglia voxel was composed of a higher fraction of grey matter.

The data collected in the BG strongly suggest that sleep loss indeed affects glutamatergic signaling on different levels. More specifically, prolonged wakefulness increased Glu, GLX and mGluR5 availability in sub-regions of the basal ganglia, and these changes were renormalized after recovery sleep. The findings corroborate and expand previously published observations from our group, showing that mGluR5 availability was increased after sleep deprivation (Hefti et al., 2013). Importantly, the new data demonstrate that recovery sleep is associated with reduced mGluR5 availability, supporting a restorative role for sleep and providing complementary evidence for the mGluR5 signaling cascade to contribute to sleep-wake regulation. The investigation of different brain regions indicated that the basal ganglia are a brain structure that reliably shows sleep-wake related changes in the glutamatergic balance in humans. The dorsal (caudate nucleus and putamen) and ventral (nucleus accumbens and olfactory tubercle) parts of the striatum and the amygdala showed increased mGluR5 availability after sleep loss (Hefti et al., 2013; supplementary material). The data strengthen the emerging hypothesis that the basal ganglia are a key player in sleep-wake regulation (Lazarus et al., 2013; Holst & Landolt, 2015; Holst & Landolt, 2018). Whereas each, the increase in Glu levels and mGluR5 availability after extended wakefulness equaled roughly 5-10 % and may be considered as small or moderate, the simultaneous changes could mutually amplify each other and cause a substantial increase in glutamatergic signaling after sleep deprivation.

### Sleep deprivation impacts on the expression of FMRP

Currently the most specific molecular marker of sleep need is the immediate early gene Homer1a (Maret et al., 2007; Mackiewicz et al., 2008) which uncouples mGluR5 from its downstream signaling partners, leading to synaptic long-term depression (Kammermeier and Worley, 2007; Ménard and Quirion, 2012; Berridge, 2016; Diering et al., 2017; Ronesi and Huber, 2008). This form of synaptic plasticity may ultimately support sleep dependent recovery processes (Diering et al., 2017; Krueger et al., 2013; de Vivo et al., 2017). The mGluR5 has been specifically associated with two proteins that may be important for sleep-wake regulation: FMRP and BDNF. Consistent with experiments in *Drosophila* (Bushey et al., 2009), we found elevated FMRP levels after prolonged wakefulness when compared to baseline. A prolonged effect of sleep deprivation might explain the delayed normalization after recovery sleep. In contrast to the findings *in vivo*, the FMRP concentration in cultured neural cells of sleep deprived rats appeared to decrease with sleep deprivation (Kwon et al., 2015). Thus, further research is needed to clarify the potential role for FMRP in sleep-wake regulation. Similarly, the evidence for a suggested role of BDNF in regulating sleep homeostasis and LTP-like plasticity after sleep deprivation (Faraguna et al., 2008; Kuhn et al., 2016) has been equivocal. Here, neither sleep deprivation nor recovery sleep revealed consistent effects on BDNF levels in the human serum as quantified with ELISA. The establishment of a reliable method to assess blood serum BDNF still remains a clinical challenge. The discrepancies among the available studies may reflect the methodological difficulties in the reliable quantification of BDNF serum concentration.

Taken together, our study provides convergent evidence that sleep deprivation and recovery sleep affects glutamatergic signaling in distinct regions of the human brain that play an important role in sleep-wake regulation. Nevertheless, the questions remain whether the observed molecular changes regulate the need for sleep or whether they reflect secondary changes associated with the expression of wakefulness and sleep, or both. The present findings warrant further studies to elucidate the mechanisms that link the homeostatic regulation of sleep and glutamatergic system activity in health and disease.

## Materials and Methods

To visualize the interplay of mGluR5 with its potential molecular signaling partners in sleep-wake regulation, a controlled in-lab study was designed, in which 3-Tesla PET/MR-Spectroscopy scanning and blood sampling were conducted three times, at the same circadian time in baseline, after 40 hours prolonged waking, and again following recovery sleep. mGluR5 availability was quantified with the novel PET radioligand [^18^F]PSS232 which is a non-competitive selective antagonist of mGluR5 (Sephton et al., 2014; Warnock et al., 2018). Concentrations of glutamate, the glutamate/glutamine (GLX) ratio and γ-amino-butyric acid (GABA) in basal ganglia (BG) and dorsolateral prefrontal cortex (dlPFC) were measured with dedicated PRESS/MEGAPRESS MRS sequences. Circulating levels of BDNF and FMRP in human blood were quantified with ELISA.

### Study Participants

The study protocol and all experimental procedures were approved by the ethics committee of the Canton of Zürich for research on human subjects. All subjects provided written informed consent prior to the experiments and received financial compensation for their participation, in accordance with the principles in the Declaration of Helsinki.

Thirty-one healthy men completed a within subject design 1-week sleep deprivation protocol after being screened for medical history and psychological state. All subjects were nonsmokers, in good health, had no history of neurologic or psychiatric disease and were instructed not to take any medications or consumed any illicit drugs 2 months prior to the study. Subjects were excluded if they traveled across multiple time zones or performing shift work 3 months prior to study participation. Subjects with unknown sleep disturbances, such as sleep apnea, sleep efficiency < 75% or periodic leg movements during sleep (PLMS) with an index of 5 or more per hour of sleep were excluded from participation based on polysomnographic screening in the sleep laboratory before enrolment. Table 1 summarizes lifestyle and demographic characteristics of the healthy study sample assessed by validated questionnaires.

Thirty-one healthy male participants, ten of which were between 60 to 70 years of age, completed the study protocol. Validated German translations and versions of questionnaires were used to assess lifestyle and personality traits. Caffeine consumption was calculated based on the following amounts per serving: coffee: 100mg; ceylon or green tea: 30 mg; cola drink: 40 mg (2 dL); energy drink: 80 mg (2 dL); chocolate: 50 mg (100 g). Diurnal preference: Horne-Östberg Morningsness-Eveningness Questionnaire (Horne et al., 1976); daytime sleepiness: Epworth Sleepiness Scale (Bloch et al., 1999); depression score: Beck Depression Inventory (Beck et al., 1961); personality traits: Eysenck Personality Questionnaire (Francis et al., 2006); cognitive assessment: Montreal Cognitive Assessment (Nasreddine, 2005); trait anxiety: State-Trait Anxiety Inventory (Spielberger et al., 1970); sleep quality: Pittsburgh Sleep Quality Index (Buysse et al., 1989).

### Pre-experimental Procedure and Experimental Protocol

Two weeks prior to the study, participants were required to refrain from all sources of caffeine and wear a wrist activity monitor on the non-dominant arm. During the 5 days prior to the study they were asked to abstain from alcohol intake and to maintain a regular 8-hour night-time sleep schedule, corresponding approximately to the participants’ habitual sleep times. Daily log-books and wrist actigraphy verified compliance with the pre-study instructions. Additionally, caffeine and ethanol concentrations in saliva and breath were tested upon entering the laboratory, to confirm participants’ abstinence.

Under constant supervision, all subjects completed a within-subject sleep deprivation protocol (Figure 1), consisting of an 8 hours adaptation and baseline night (time in bed: 11:00_PM_-07:00_AM_), followed by 40 hours of continuous wakefulness, and terminated by a 10 hour recovery night. In baseline, sleep deprivation and recovery condition, twenty-two subjects underwent a combined positron emission tomography (PET) and magnetic resonance spectroscopy (MRS) examination at the same circadian timepoint (4:23_PM_ ± 23 min) with [^18^F]PSS232 to quantify mGluR5 availability in the brain (Division of Nuclear Medicine, University Hospital Zürich). Due to time and logistic constraints, only two subjects could be PET scanned per experimental week. To optimize data collection, one additional subject was included in each study block (9 in total) as a back-up candidate, participating in the entire experimental protocol, MR imaging and blood sampling, but without the [^18^F]PSS232 injection and PET scan.

### Magnetic Resonance Spectroscopy data acquisition and analysis

Magnetic resonance spectroscopy data were acquired simultaneously with the PET data using a GE 3T combined PET/MR scanner (SIGNA PET/MR; GE Healthcare). Single-voxel edited ^1^H-MR spectra were acquired from two voxels of interest in the left dorsolateral prefrontal cortex (dlPFC; 30 x 25 x 40 mm^3^) and in the basal ganglia (BG; 25 x 25 x 25 mm^3^) using the Point RESolved Spectroscopy (PRESS) and MEGAPRESS methods. In addition, a third VOI in the BG (25 x 30 x 35 mm^3^) was measured with the MEGAPRESS method (Mescher et al., 1998) to specifically quantify GABA. To ensure a consistent MRS voxel position between subjects, the voxel was carefully positioned based on anatomical landmarks on the T1 image. The T1 weighted MR images were also used to correct for partial volume effects related to the CSF content in the MRS voxel, as well as for gray/white matter correction.

MEGAPRESS: A total of 320 spectra were averaged to obtain the final spectrum. Individual spectra were acquired with a TR of 1800 ms, an echo time of 68 ms, and an eight-step phase cycle, resulting in a total acquisition time of ~10 minutes. For each metabolite spectrum, 16 water reference lines were also acquired as part of the standard PROBE acquisition.

PRESS: The PRESS spectra were acquired with an echo time (TE) of 35 ms and a repetition time (TR) of 3 ms. 160 spectral averages were acquired to obtain the final spectrum resulting in an acquisition time of 9 min.

### Data analysis

MR spectra were analyzed with LCModel v. 6.3-1 (Provencher 1993), which is a fully automated spectral fitting method. For the MEGAPRESS data, edited spectra were analyzed with a simulated basis set providing metabolite concentrations for glutamine (Gln), glutamate (Glu), glutamate to glutamine (GLX), GABA, N-acetylaspartate, and glutathione. The control parameter sptype = ‘megapress-2’ was used to avoid mis-assignment of the baseline to GABA. For the PRESS spectra, a standard experimental basis set was used, from which data for creatine, glutamate to glutamine, myo-inositol, N-acetylaspartate, and total choline were extracted (Supplementary Figure S1). For all spectra, peaks that were poorly fitted, resulting in Cramer-Rao minimum variance bounds of more than 20 % as reported by LCModel, were excluded from further analyses.

### PET Image Acquisition

A T1-weighted, whole-brain, three-dimensional magnetic resonance (MR) image (resolution: 1 x 1 x 1 mm) was obtained for each subject in parallel to the PET imaging (SIGNA PET/MR 3T whole-body PET/MR unit equipped with an 8-channel head coil; GE Healthcare), to exclude morphological abnormalities and as anatomical standard for the quantification of the PET images. After an automated standard single bolus injection of [^18^F]PSS232, dynamic PET brain imaging was performed for 60 min. Images were acquired in 3D Mode with Time of flight fully iterative reconstruction (VPFX) using standard MRAC based attenuation correction with a resolution of 1.17 x 1.17 x 2.78 mm^3^ and Matrix size of 256 x 256 x 89 voxels binned into 43 timeframes (11 x 1 min, 22 x 2 min, 10 x 1 min). Subjects were instructed to not fall asleep during image acquisition. To verify wakefulness, subjects were instructed to gently press the button of a response box, generating as little movements as possible. As soon as subjects stopped pressing the response box, subjects were alerted via an intercom. Direct contact was avoided, to minimize movement artifacts.

Injected activity (baseline: 164.7 ± 24.5 MBq; sleep deprivation: 159.1 ± 15.9 MBq; recovery: 154.7 ± 12.9 MBq) did not differ between the conditions (p_all_ > 0.45; two-tailed, paired t tests).

### Image processing and quantification

All processing and quantification analyses were conducted with a dedicated brain PET/MR analysis tool (PNEURO, version 3.7) provided by PMOD Technologies LLC. PET image processing consisted of within-subject rigid-body motion correction followed by time-series alignment to the MR-T1 image for between scan comparisons. For PET quantification, the T1 image was automatically segmented, separating the MR image into gray matter (GM), white matter (WM) and cerebrospinal fluid (CSF) probability maps. After matching the T1 MR image to the functional PET images, the specific neocortical and subcortical (core brain segments) brain regions were determined using the Hammers-N30R83 brain atlas. Partial volume correction (PVC) was performed automatically in the PNEURO toolbox. A time activity curve (TAC) was calculated for each VOI. Because a single bolus injection was used, the binding potential (BPnd) was quantified with standard SRTM2 [Simplified Reference Tissue Model with fixed k2; (Wu & Carson, 2002)] modelling. For modelling, TACs of receptor-rich regions (gray matter VOIs) were compared to the TAC of a receptor-less region (cerebellum) believed mainly to entail non-specific binding (Warnock et al., 2018).

### Assessment of proteins from human serum

Fresh blood was collected immediately before the PET/MRS scans in two 10ml cloth activator tubes (BD Vacutainer^®^ CAT). The samples were allowed to cloth for about 30 minutes at room temperature (RT) before centrifugation (2.000 relative centrifugal force (RCF) for 10 min). 1.9 mL serum was extracted and purified by a second centrifugation step (12.000 RCF for 5 minutes). The purified serum was aliquoted into multiple 255μ1 samples and stored in Eppendorf tubes (SafeSeal micro tube 1.5ml, PP, Sarstedt, Nümbrecht) The probes were then snap-frozen in liquid nitrogen and stored at -80°C for future analysis.

### Fragile Xmental retardation protein (FMRP)

FMRP was studied by a quantitative sandwich enzyme-linked immunosorbent assay (ELISA) purchased prefabricated and ready to use (Human Fragile X mental retardation 1 ELISA kit, MyBioSource, San Diego, California USA). The detection rate of this assay is 15.6-1000 pg/ml. A 96-well microplate was pre-coated with a FMRP-specific antibody. Each sample was quantified at least twice for independent confirmation. The assay was performed according to the manufacturer’s instructions and guidelines.

### Brain-derived neurotrophic factor (BDNF)

Quantification of serum BDNF levels was conducted at the Department of Clinical Psychology and Psychotherapy at the University of Zurich using a 96-Well MULTIARRAY^®^ BDNF Assay purchased from Meso-Scale Discovery (MSD^®^, Rockville, Maryland USA). The analysis was performed according to the manufacturer’s instruction.

### Statistical Analyses

All statistical analyses were performed with SAS 9.4 software (SAS Institute, Cary, North Carolina). If not stated otherwise, numbers represent mean ± standard error of the mean (SEM). Only significant results are reported. Following standards, the error bars shown in the figures represent the SEM of between-subjects variability. Mixed-effect repeated measure analysis of variance included the factors ‘condition’ (baseline, sleep deprivation, recovery). Analyses of PET data were limited to the predefined VOIs and strictly statistically corrected as follows: 1. Comparisons of the mixed effect factor ‘condition’ for each VOI were post-hoc corrected (Tukey-Kramer correction: α < .05). 2. Corrected Tukey-Kramer p-values across all investigated VOIs were additionally corrected using false discovery rate correction [(FDR): α < .05]. Following significant main effects or interactions, Mann-Whitney *U* testing was employed to illustrate individual differences.

## Acknowledgements

This work was supported by the Swiss National Science Foundation grant # 320030_163439 to HPL. We thank I. Clark, D.M. Baur, S.M. Pereira Soares, A. Dieffenbacher and S. Brühlmeier for their help with data collection and analyses. Furthermore, we thank S. Geistlich for the help and production of the radioactive ligand and Prof. Dr. M. Kohler as well as Prof. Dr. S. Brown for providing access to their laboratories for blood processing and analyses.

## Author contributions

**Susanne Weigend**, Data curation, Formal analysis, Supervision, Investigation, Visualization, Writing-original draft, Writing-review and editing; **Sebastian C Holst**, Data curation, Formal analysis, Supervision, Funding acquisition, Investigation, Visualization, Writing-original draft, Writing-review and editing; **Valerie Treyer**, Resources, Methodology, Supervision, Writing-review and editing; **Ruth L. O’Gorman Tuura**, Software, Methodology, Data curation, Writing-review and editing; **Josefine Meier**, Data curation, Formal analysis, Investigation; **Simon M. Ametamey, Alfred Buck**, Resources, Methodology, Project administration; **Hans-Peter Landolt**, Conceptualization, Resources, Data curation, Supervision, Funding Acquisition, Writing-original draft, Project administration, Writing-review and editing.

---

**Supplementary Table S1:** Behavioral Effects of Sleep Deprivation

|  | Baseline (BL) | Sleep deprivation<br>(SD) | Recovery<br>(RE) | SD vs. BL<br>*p* < | RE vs. SD<br>*p* < | RE vs. BL<br>*p* < |
| --- | --- | --- | --- | --- | --- | --- |
| Karolinska Sleepiness Scale | 3.23 ± 0.19 | 4.92 ± 0.32 | 2.81 ± 0.19 | **<.001** | **<.001** | **<.013** |
| Tiredness Symptoms Scale | 0.52 ± 0.12 | 1.59 ± 0.29 | 0.43 ± 0.15 | **<.001** | **<.001** | <.609 |
| Visual Analogue Scales of States |  |  |  |  |  |  |
| Mood | 66.55 ± 1.90 | 61.69 ± 2.20 | 70.98 ± 2.05 | **<.001** | **<.001** | **<.001** |
| Energy | 63.48 ± 1.64 | 55.07 ± 2.11 | 65.59 ± 2.22 | **<.001** | **<.001** | <.240 |
| Motivation | 32.81 ± 2.00 | 38.14 ± 2.36 | 27.45 ± 2.13 | .066 | **<.001** | **<.048** |
| Agitation | 41.43 ± 3.38 | 35.09 ± 2.56 | 33.88 ± 2.98 | .073 | .561 | **.006** |

Values represent means ± SEM (n = 31). *P* values refer to two-tailed, paired *t* tests. Significant differences between conditions are highlighted in bold.

**Figure S2:**
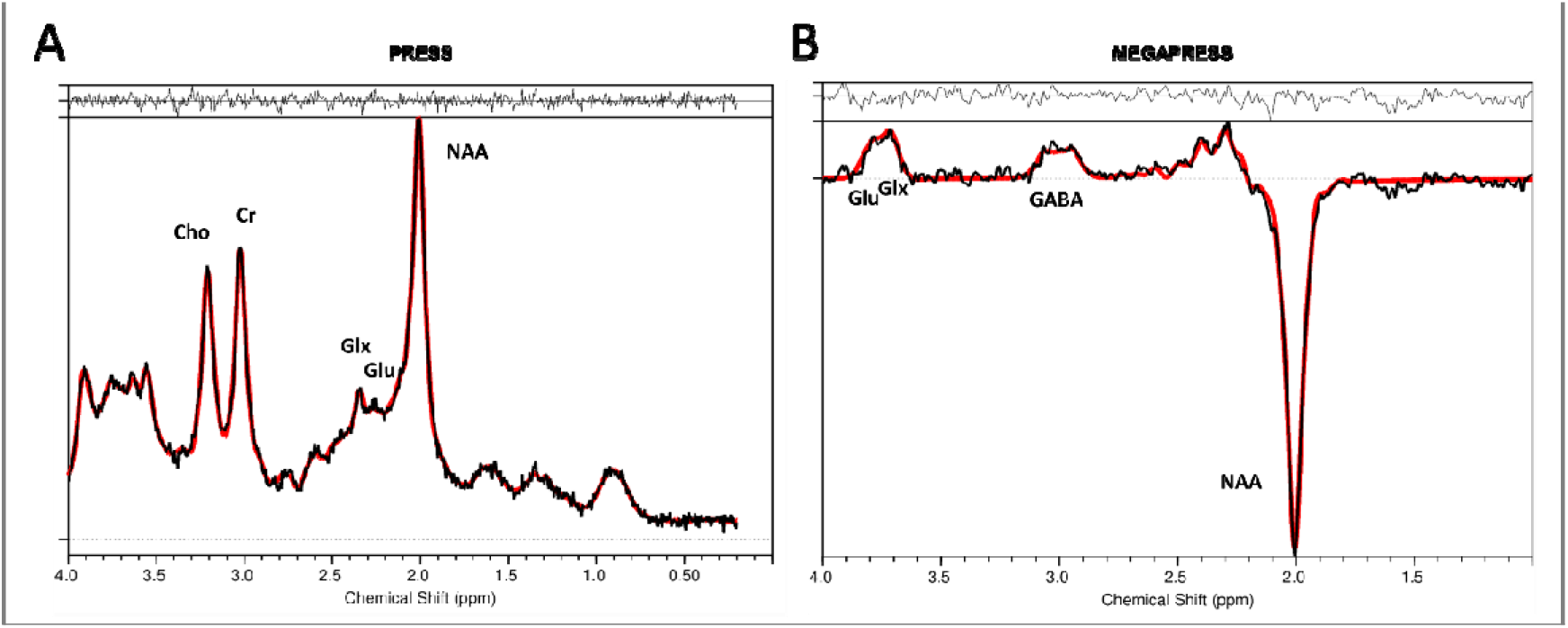
Example data of MRS PRESS and MEGAPRESS spectra. Representative PRESS (left panel) and MEGAPRESS (right panel) spectra. Each spectrum is plotted in black with the LCModel fit overlaid in red. The residuals of the fit are also plotted above each spectrum.

**Figure S1:**
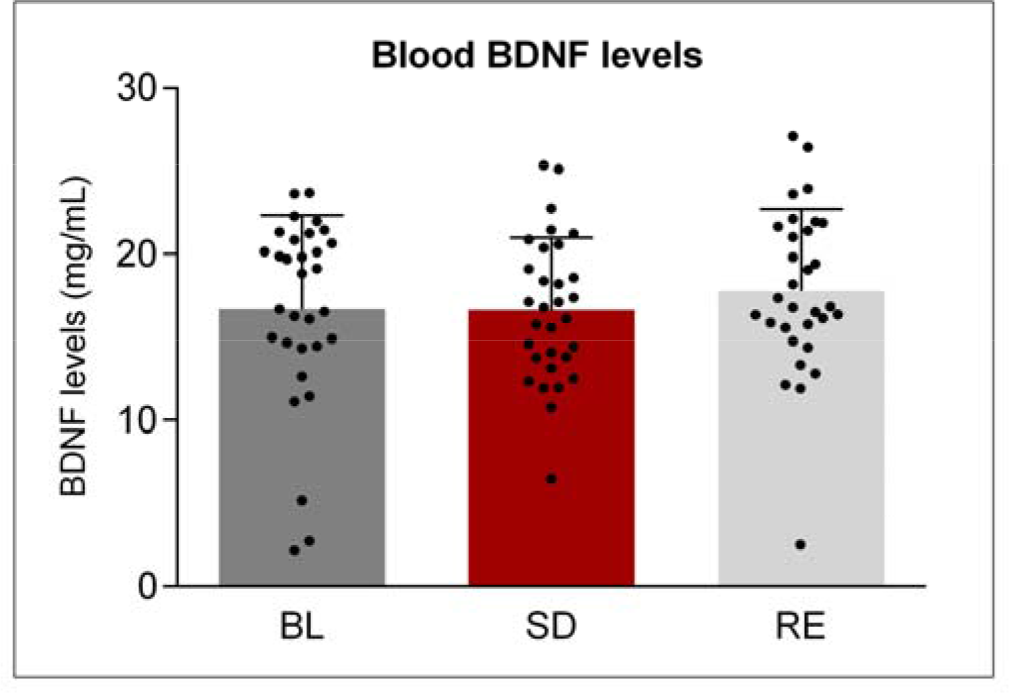
Effects of sleep deprivation and recovery sleep on circulating blood brain derived neurotrophic factor protein levels. Columns display blood BDNF levels in baseline (BL, dark grey), sleep deprivation (SD, red) and recovery (RE, light grey) conditions. Data represent means + standard error of the mean (SEM) in N=22. Black dots represent individual subjects. Mixed model ANOVA main effect of ‘condition’: F_2,90_ = 0.23, p > 0.7.

